# A Diffusion-Based Autoencoder for Learning Patient-Level Representations from Single-Cell Data

**DOI:** 10.1101/2025.08.21.671613

**Authors:** Rebecca Boiarsky, Johann Wenckstern, Nicholas J. Haradhvala, Gad Getz, David Sontag

## Abstract

Single-cell RNA sequencing (scRNA-seq) offers insights into cellular heterogeneity and tissue composition, yet leveraging this data for patient-level clinical predictions remains challenging due to the set-structured nature of single-cell data, as well as the scarcity of labeled samples. To address these challenges, we introduce scSet, a diffusion-based autoencoder that learns patient-level representations from sets of single-cell transcriptomes. Our method uses a transformer-based encoder to process variably sized and unordered cell inputs, coupled with a conditional diffusion decoder for self-supervised learning on unlabeled data. By pre-training on large-scale unlabeled datasets, scSet generates robust patient representations that can be fine-tuned for downstream clinical prediction tasks. We demonstrate the effectiveness of scSet patient embeddings for clinical prediction across multiple real-world datasets, where they outperform existing patient representations, even with limited labeled data. This work represents an important step toward bridging the gap between single-cell resolution and patient-level insights. Code is available at https://github.com/clinicalml/scset.

## 1. Introduction

Single-cell RNA sequencing (scRNA-seq) provides a detailed view of the cellular composition of a tissue, enabling insights such as the identification of cell type specific biomarkers (4; 17), tumor heterogeneity (1; 4), the composition of the tumor-immune microenvironment (43; 35), and the diverse cell states that can exist for a single-cell type (31; 38). While it is widely acknowledged that single-cell features correlate with clinical outcomes of interest (32), few machine learning (ML) tools exist to make patient-level predictions based on scRNA-seq data. We identify two major hurdles for ML for patient-level prediction from single-cell data: (1) Patient single-cell datasets do not immediately lend themselves to predictive machine learning methods which assume that the input features are either consistently or meaningfully ordered (e.g. logistic regression or recurrent/convolutional networks, respectively), since their features (the cells) have no consistent or meaningful order. (2) The numbers of single-cell samples with clinical labels for any given disease state is often too small to employ machine learning methods to automate the discovery of features which correlate with clinical outcomes of interest. This is due to a combination of single-cell data being expensive to generate, clinical samples requiring patient consent and potentially invasive biopsies, and clinical labels requiring careful annotation and patient follow-up.

Most commonly, single-cell samples are simply averaged across cells prior to being input to an ML model, or else manual feature engineering and statistical analysis are used to find correlations between single-cell features with patient-level information (33; 44). Machine learning architectures that predict sample-level information from a parametrized embedding of single-cell data have only recently started to emerge (11; 21; 23).

In this work, we introduce scSet, a diffusion-based autoencoder for learning patient-level representations from scRNA-seq data. Our method addresses both of the above challenges, first by developing an encoder that can handle variably sized and unordered cell inputs, and second by leveraging unlabeled samples for self-supervised learning via a denoising diffusion objective. Taken together, scSet learns patient representations in an unsupervised manner, which can then be fine-tuned to predict clinical features of interest from a limited cohort of clinically-labeled samples.

## 2. Related Work

### 2.1. Representation Learning for Sets

Representation learning for sets has been explored through various approaches, both supervised and unsupervised, with applications spanning multiple domains, including point clouds, graphs, and multi-instance learning (MIL). Early work such as DeepSets (42) proposed permutation-invariant architectures that aggregate unordered inputs using pooling functions like sum or mean. Transformer-based models for sets, such as Set Transformer (19) and Attention-based Deep Multiple Instance Learning (16), introduced attention mechanisms to capture higher-order interactions between set elements. More recently, unsupervised approaches such as SetVAE (18) incorporate principles from Set Transformer into a variational autoencoder framework to learn unsupervised latent representations of sets, using the Chamfer Distance as a proxy reconstruction loss in order to handle unordered set data. A noise prediction loss as used in diffusion models (15; 28) is less computationally expensive and can also handle unordered set data, and thus we were motivated to explore a diffusion-based decoder for unsupervised set learning. Others have recently begun to explore this direction too, with applications in 3D point clouds instead of biology (45). Our work builds on these foundational principles but extends them to the biomedical domain, enabling unsupervised patient representation learning from sets of single cells.

### 2.2. Representation Learning for scRNA-seq

Representation learning in the single-cell space has mostly focused on learning representations of individual cells, with methods such as scVI (24), Geneformer (37), and scGPT (7), as well as multimodal models such as totalVI (12). Any of these cell embeddings can be used as input to our model, which instead focuses on encoding a set of cells into a patient-level representation, and decoding a patient representation back to individual cells through conditional denoising diffusion. Other approaches for learning sample-level encodings of single-cells have only recently started to emerge and have focused on supervised methods, as described in Section 2.3.

### 2.3. Patient-Level Representations from scRNA-seq

The simplest and most common method for summarizing sample expression is to take the average gene expression across all cells in the sample, referred to as pseudobulk. However, pseudobulk obscures the granular view of cell states afforded by single-cell. Recently, a few methods have been proposed for learning patient-level representations from scRNA-seq data (21; 11; 23; 8; 26; 13; 9; 40). These have mostly built off of the deep set (42) or attention-based multiple instance learning (ABMIL) (16) frameworks. While these works propose architectures that learn to aggregate single-cell data into patient-level representations, they are all trained on (semi-)supervised tasks. By contrast, a key contribution of our work is our self-supervised training objective, which allows learning representations from unlabeled data and improves the quality of our downstream supervised predictions.

### 2.4. Diffusion Models for scRNA-seq data

Diffusion models have been used for a variety of tasks in machine learning, ranging from image generation (15) to drug discovery (6). Recently, diffusion models have been applied to scRNA-seq data for gene expression imputation (25; 22; 10). While these methods use diffusion models to generate scRNA-seq data, they do not condition the model on a patient-specific representation, or leverage it as part of an autoencoding framework for learning a patient representation.

## 3. Method

Single-cell RNA sequencing profiles the transcriptomes of individual cells in a patient sample. Each cell’s transcriptome is represented as a vector of gene expression values, *x* ∈ ℝ^*G*^, where *G* denotes the total number of all genes detected across cells in our dataset. A scRNA-seq sample from a given patient is observed as an unordered set of single-cell transcriptomes, 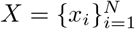. Our goal is to learn a mean-ingful vector representation *z* ∈ ℝ^*d*^ of the set of cells for each patient.

To this end, we propose scSet, a diffusion-based autoencoding framework for learning patient-level representations from scRNA-seq data. The following sections detail the decoder, encoder, and training procedure for scSet. A schematic overview of the model is provided in Figure 1.

**Figure 1.**
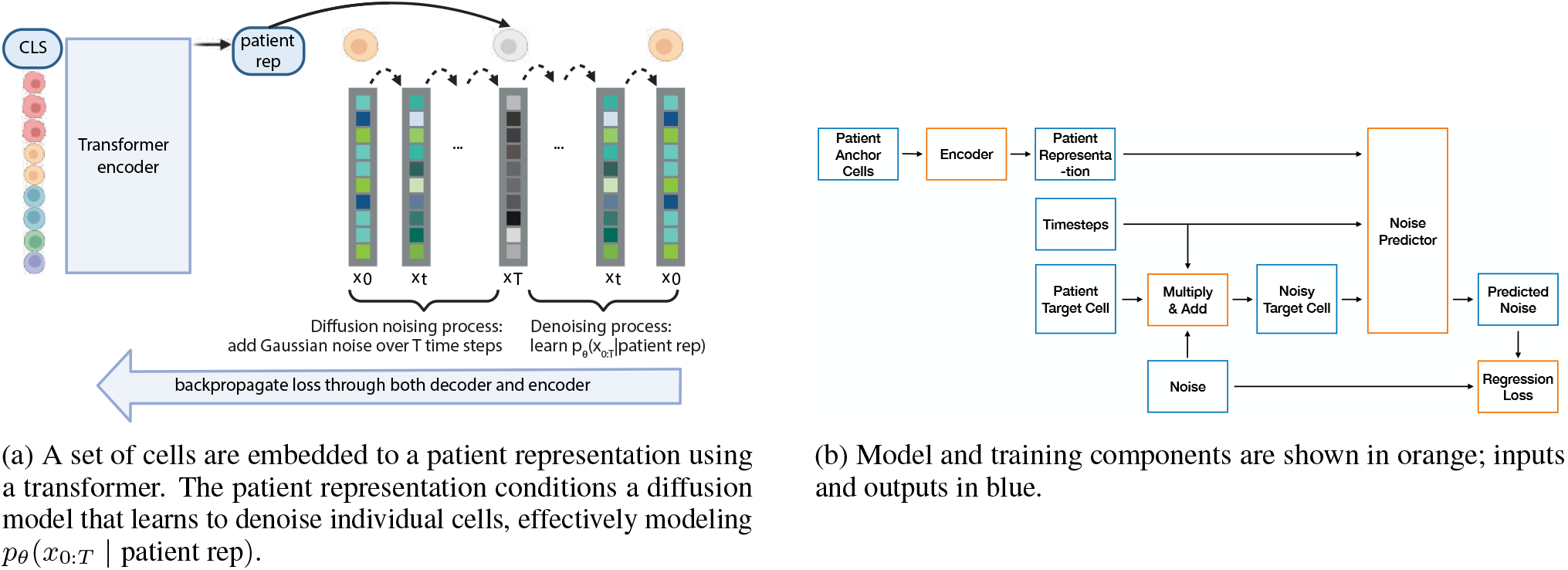
Two complementary overviews of the SCSet model: (a) schematic of the patient-conditioned diffusion model, and (b) model and training flowchart.

### 3.1. Diffusion-based Decoder

We employ a conditional Denoising Diffusion Probabilistic Model (DDPM) (15; 28), which uses the patient representation *z* for conditioning and, starting from noise, generates sample cells matching the patient profile.

Given a number of time steps *T* ∈ ℕ and a variance schedule *β*_1_, …, *β*_*T*_ ℝ_*>*0_, we model the diffusion forward-process as

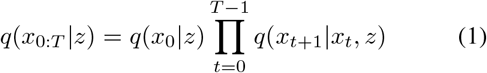

where *q*(*x*_0_|*z*) is the data distribution of cell profiles conditioned on a patient representation *z* and

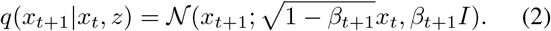

As a decoder, we learn a denoising backward-process

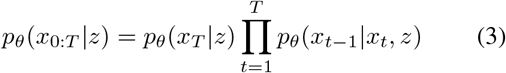

parametrized as

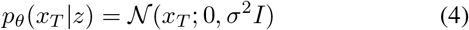

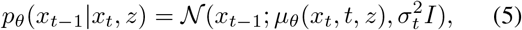

with mean predictor *µ*_*θ*_(*x*_*t*_, *t, z*) given by

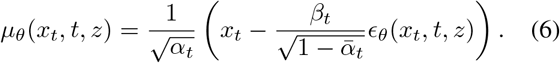

Here *ϵ*_*θ*_(*x*_*t*_, *t*) is a noise-predicting neural network, *α*_*t*_ = 1 − *β*_*t*_ and 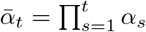.

A key assumption of our paper is that cells in a sample are conditionally independent given *z*, and thus the probability of the set of cells *X* is the product of the probability of each cell in the set, 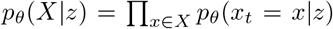. Due the presence of multiple cell types in each sample, we expect *p*_*θ*_(*x*_*t*_ = *x*|*z*) to be a complex, multimodal distribution.

The noise-predicting network is an adapted multilayer perceptron with residual connections. The patient representations and sinusoidal time step embeddings are each processed by feed-forward networks, summed and incorporated into the noise prediction through Adaptive Layer Normalization (29). For the diffusion process, we use a cosine noise schedule (28) and *T* = 1000 time steps.

### 3.2. Transformer-based Encoder

The scSet encoder, *f*_*ϕ*_, maps the unordered set of cells in a patient sample 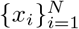 to a fixed-dimensional representa-tion *z ∈* ℝ^*d*^, where *d* = 256 in our model. To address the challenges posed by the variable size and lack of ordering in the input data, we employ a transformer-based architecture with a learnable [CLS] token that serves as a global representation of the input set.

The architecture begins with a linear embedding layer that projects each cell *x*_*i*_ *∈* ℝ^*G*^ into *d*-dimensional space:

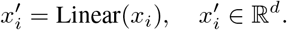

A learnable [CLS] token is appended to the set of cells, forming the input to the encoder:

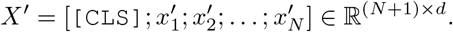

The transformed set *X*^*′*^ is then passed through a series of transformer encoder blocks (39), each consisting of multi-head self-attention, feedforward networks, and layer normalization. These layers are intended to model interactions between cells and to encode information about cells in the context of their tissue environment. Formally, each encoder layer is defined as:

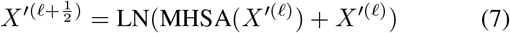

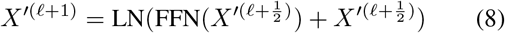

where MHSA denotes multi-head self-attention, LN Layer Normalization and FFN a feed-forward network. Dropout is applied to attention and feed-forward layers to prevent overfitting.

After passing through *L* transformer blocks, the embedded [CLS] token is extracted as the patient representation *z*. This representation *z* serves as the input to the diffusion-based decoder for patient-specific cell generation, and the encoder is jointly optimized with the decoder during training, as described in Section 3.3.

When using *z* as input for a downstream clinical prediction task, the weights of the encoder can optionally be further updated to tailor the patient embedding for the downstream task (see Section 4.2.3).

### 3.3. Training Procedure

The encoder *f*_*ϕ*_ and the decoder *p*_*θ*_ are jointly trained using the noise prediction loss

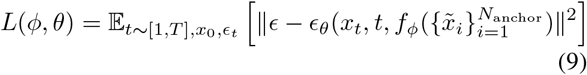

where *t* ∼𝒰(1, …, *T*), *ϵ* ∼𝒩(0, *I*), *x*_*t*_ ∼ *q*(*x*_*t*_ *x*_0_, *z*), and *x*_0_ as well as 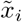 for *i* ∈ 1, …, *N*_anchor_ are drawn uniformly without replacement from the cells of patient *j*. To estimate the loss during training in practice, mutually exclusive subsets of *X* are used as input to the encoder (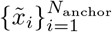, referred to as “anchor cells”) and to the noise-prediction network (*x*_0_, referred to as a “target cell”). Note that this objective functions allows back-propagation of the gradients through both the denoising decoder *p*_*θ*_ and the sample encoder *f*_*ϕ*_.

The training procedure without mini-batching is described in detail in Algorithm 1. For training, we use the AdamW optimizer with learning rate 1 × 10^−3^. Gradients are clipped at a threshold of 0.1.

## 4. Experiments

We evaluate scSet’s learned patient embeddings through (i) qualitative and quantitative evaluations of the unsupervised trained embeddings and (ii) using the patient embeddings as input to downstream clinical prediction tasks. We describe the datasets, metrics, baselines and results for each of these approaches in Sections 4.1 and 4.2, respectively.

### 4.1. Training Patient Representations via Conditional Diffusion

In this section, we validate that the patient representations learned by scSet capture known variations between patients. First, we show that the diffusion model decodes the expected distribution of cell types for each patient, and then we use real and semi-synthetic data to validate that patients with known differences are separated in the latent space.

#### 4.1.1. Data

The scSet autoencoder was trained on data from the CZ CELLxGENE (CxG) Discover Census (3), containing 7,342 samples from diverse tissue and disease contexts. scVI embeddings provided in the Census were used to represent input cells; we retained 14 scVI latents by filtering to latents with standard deviation *>*.4 across the pretraining corpus. We chose to represent cells using scVI embeddings rather than raw gene expression as it already partially corrects for batch effects and reduces the dimensionality of inputs. We used 90% of patients for training and held out 10% for evaluation.

For our semi-synthetic data, we created patient samples by resampling cells from a pool of 8064 immune cells (natural killer cells, helper T cells, CD8+ cytotoxic T cells, and monocytes) from 32 patients from a multiple myeloma study (43) that was not part of the pretraining corpus, in order to create synthetic patients belonging to different synthetic “patient subtypes.” For each subtyping experiment, we simulated 12 patients, each with 200 cells. For our cell type composition experiment, we created samples that were enriched for a given cell type: we randomly sampled cells of the dominant cell type to account for 55% of the sample composition, and the remaining cell types to each account for 15%. For our perturbation subtyping experiment, we created a “perturbed” subtype in which cell types were present in equal proportions to their unperturbed counterparts, but with a slight phenotypic shift in helper T cells, and an equal and opposite shift in CD8+ cytotoxic T cells (more details provided in Appendix G).

#### 4.1.2. Results

If the model has learned meaningful patient representations, we expect the diffusion decoder—which is conditioned on those representations—to generate sets of cells that closely resemble a patient’s true cells. Thus, we ran inference on the CxG evaluation set (10% of patients that were not used to train scSet), decoding 500 cells per patient (the number of decoded cells is arbitrarily set by the user). Starting from Gaussian noise at time step *T* = 1000, we visualize via UMAP (27) the reconstructed cell profiles of patients grouped by tissue as they are denoised over time steps in the diffusion decoder (Figure 2). Color-coding cells by their cell types shows that for each tissue, the model generates the same cell types which were present in the ground truth single-cell data, and in relatively similar proportions (the Pearson correlation coefficient between the true and reconstructed cell type proportions for each tissue was consistently high: 0.95 for lung, 0.97 for breast, 0.89 for heart, and 0.91 for blood). We include tables showing the true and reconstructed cell type proportions for each tissue in Appendix H. Note that since generated cells have no ground truth cell type labels, we predicted instead pseudo-cell type labels from their simulated profiles at *t* = 0 using a *k*-Nearest Neighbors classifier trained on the true cells from these patients.

**Figure 2.**
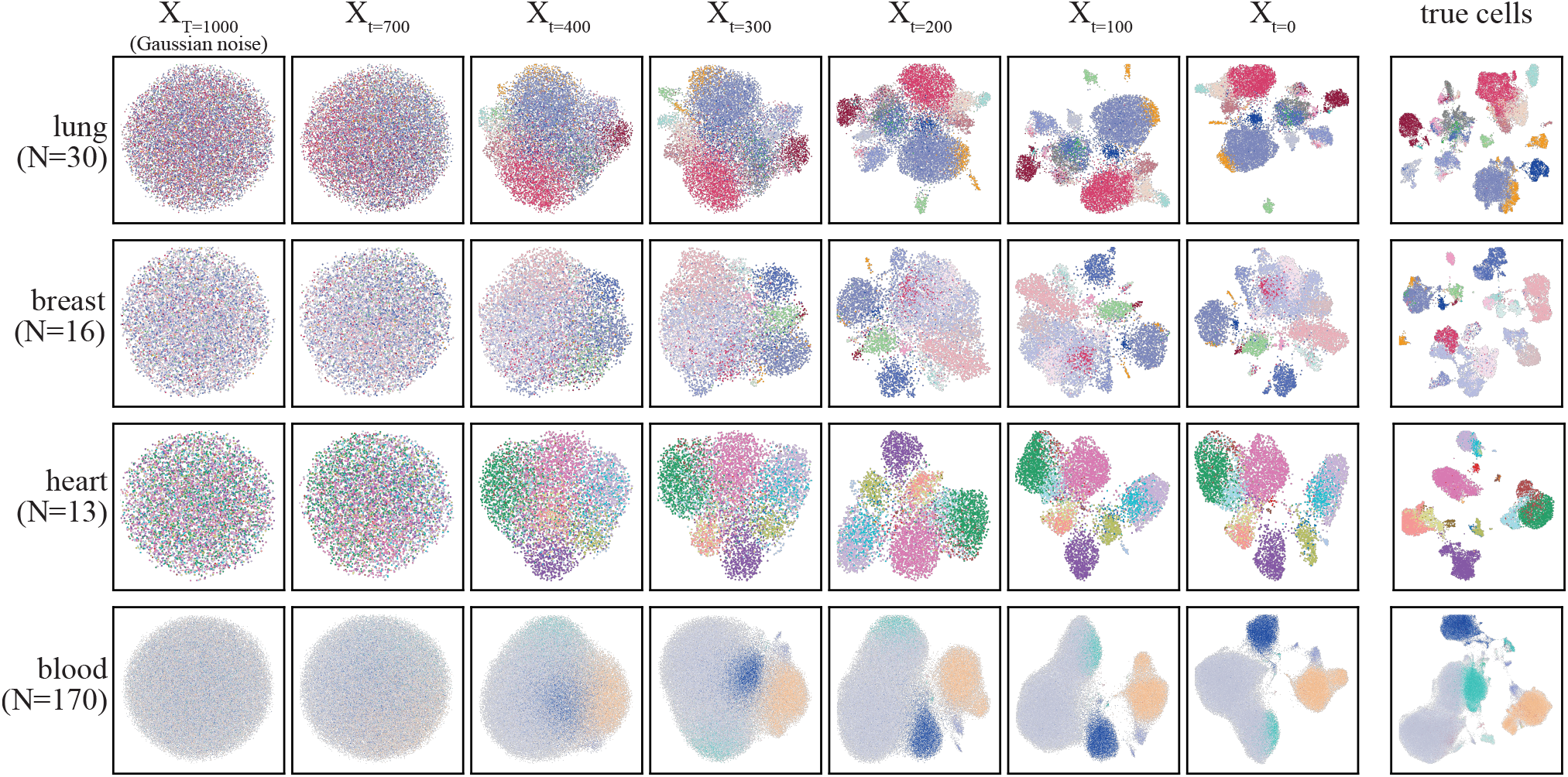
UMAP visualizations of cells over timesteps of the denoising process, starting from random noise (*X*_*T* =1000_). Test set patient samples were embedded using *f*_*ϕ*_ and used to condition the denoising network; the union of these cells is shown in the right column, labeled “true cells.” 500 cells per patient were generated. For visualization purposes, each row contains the union of cells from a given tissue, with the number of patients *N* indicated in parentheses. True cells are colored by their ground-truth cell type and simulated cells are colored by pseudo-cell type labels, obtained by predicting cell types using a *k*-Nearest Neighbors classifier trained on all cells in the test set.

While Figure 2 and Appendix H show that the landscape of true cells for each tissue matches the landscape of generated cells, we wanted to confirm that the patient representations condition the model to generate *patient-specific* profiles, rather than simply tissue-specific profiles. We calculated the Pearson correlations between the cell type proportions vectors of the true and generated cells for each sample in a tissue, and observed that the generated cell type distribution for a given patient is usually most strongly correlated with the ground-truth distribution of cell types in the patient used for conditioning. We visualize these results in Figure 3.

**Figure 3.**
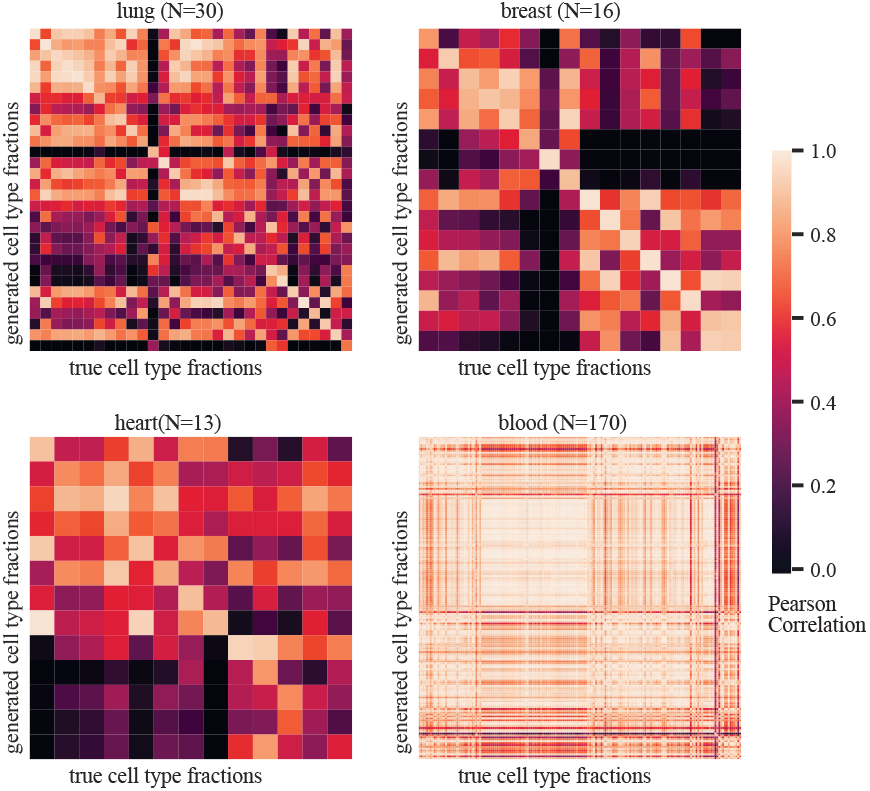
Pearson correlations between true and reconstructed cell type proportions for evaluation set patients from each tissue (the same patients shown in Figure 2). scSet reconstructs a set of cells per patient which matches the relative proportions of cell types in the true patient sample.

Finally, we qualitatively validate that the patient representations learned are reasonable. First, we inspect the embeddings for 1,500 patient samples from the CxG Discover Census. Coloring each patient sample by its tissue type, Figure 4 reveals that scSet representations separate patient samples by their tissue origin, as expected. We next ran hierarchical clustering on patient embeddings from our semi-synthetic data, and observed that scSet separates patients based on differences in cell type proportions (Figure 5a) as well as shifts in cell states, or phenotypes (Figure 5b). For the perturbation experiment, we intentionally induced a perturbation that could not be detected between samples whose cells had simply been averaged (since equal and opposite perturbations were imposed on different cell types), highlighting that scSet captures signal in its patient embeddings that pseudobulk would not be able to (Figure 5c).

**Figure 4.**
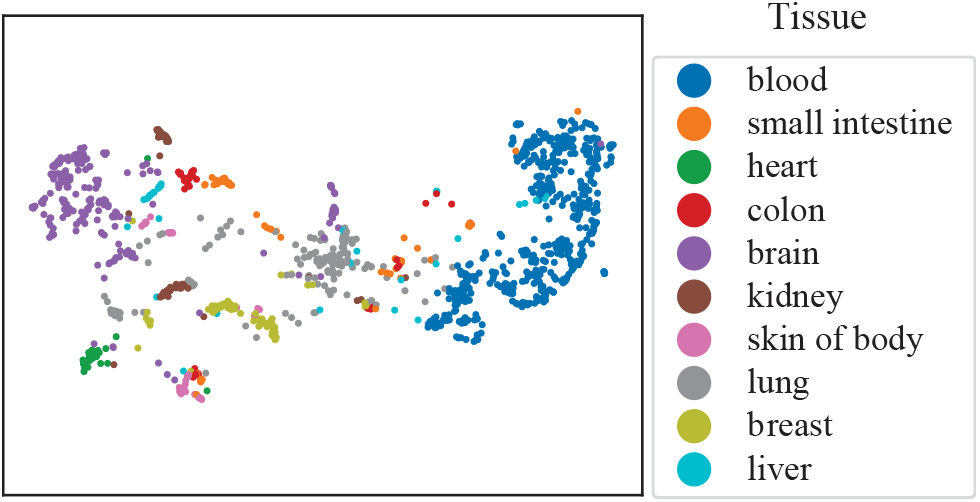
UMAP visualization of the patient embeddings encoded via scSet, colored by tissue type.

**Figure 5.**
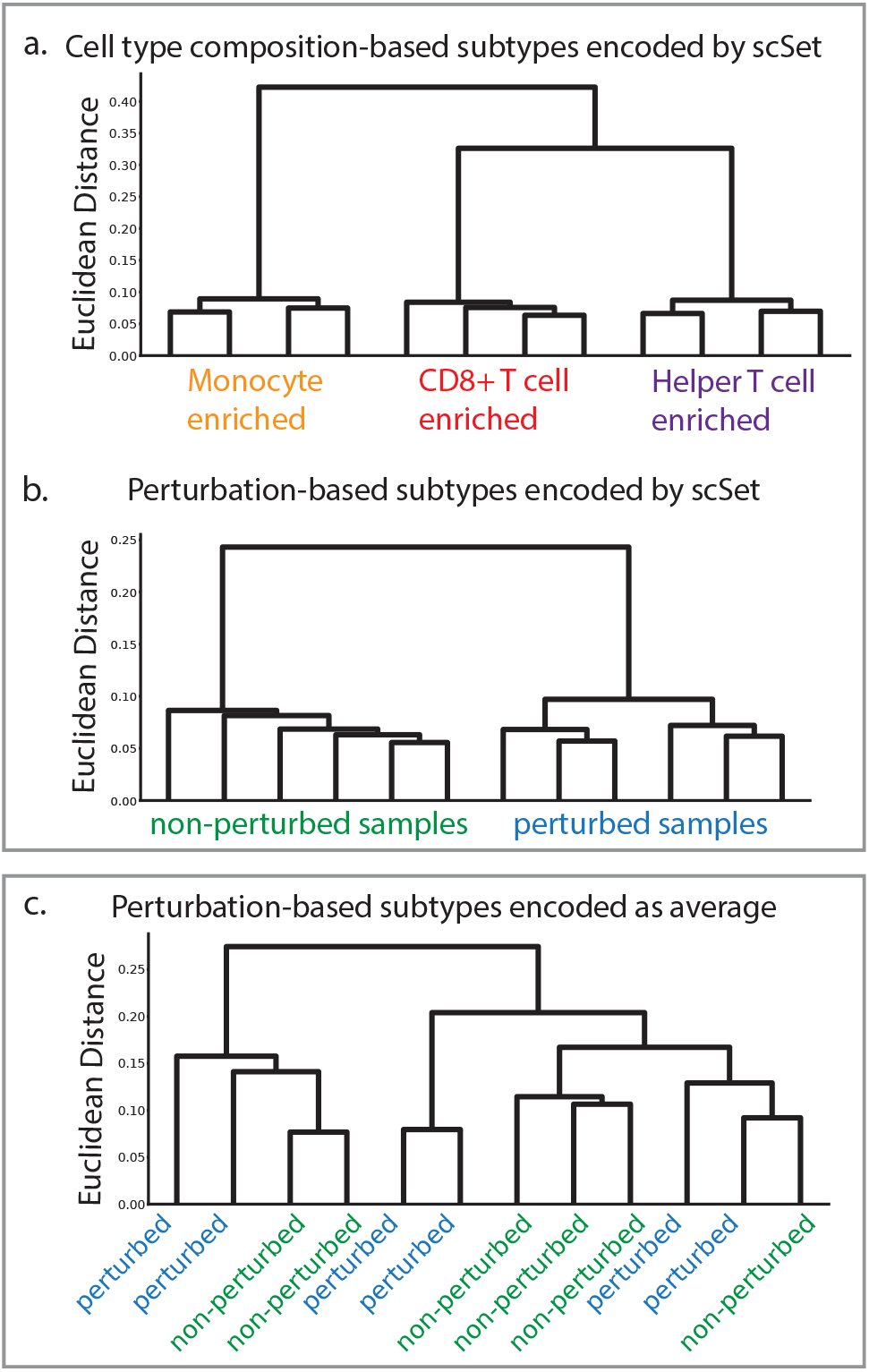
Hierarchical clustering of semi-synthetic samples that were generated as part of the (a) cell type composition subtype experiment or (b) cell type perturbation subtype experiment. The left box shows that scSet embeddings for these semi-synthetic patients cluster by subtype. The right box (c) shows that a simple Average embedding of the perturbed patients do not cluster by subtype.

### 4.2. Clinical Prediction from Patient Representations

We evaluated our learned embeddings by using them as input to supervised models for predicting patient-level phenotypes for the datasets described below in Section 4.2.1. Our clinical prediction models are comprised of scSet’s transformer encoder, which aggregates single-cell data into a patient-level representation via the [CLS] token, and an appended prediction head, as described in Section 4.2.3.

#### 4.2.1. Data & Tasks

##### HLCA

The human lung cell atlas (HLCA) (34) combines 49 datasets related to the human respiratory system, integrating over 2.4 million cells from 486 individuals. In our *HLCA triple* task, we train models to discriminate between the three most prevalent disease states for lung tissue samples in this dataset: normal (*N* = 216 samples), COVID-19 (*N* = 82), and pulmonary fibrosis (PF) (*N* = 71). We use a 10-fold cross validation scheme and assign all the patients from a given dataset to the same fold to avoid confounding by dataset-specific batch effects. This setup requires the model to generalize across batches, and is significantly more challenging than an ungrouped K-fold scheme, but better reflects a potential real-world deployment setting for our model. We include results from a binary version of this task, discriminating between normal and PF patients, in Appendix I.

##### SLE

The systemic lupus erythematosus (SLE) dataset (30) contains 1.2 million peripheral blood mononuclear cells (PBMCs) from 162 patients with SLE and 99 healthy controls. We train models to discriminate between SLE and healthy samples, and use a standard 10-fold cross-validation scheme for evaluation.

##### COVID-19

The COVID-19 dataset (36) profiles transcriptomes of 624,325 peripheral blood mononuclear cells from 24 healthy donors and 102 patients with varying severities of COVID-19, ranging from asymptomatic to critical disease. We train models to discriminate between COVID-19 and healthy samples, and use a standard 10-fold cross-validation scheme for evaluation.

Each dataset was pre-processed to embed cells using the trained scVI model available on CxG Discover Census. The same 14 latent dimensions as used for pretraining scSet were retained.

#### 4.2.2. Baseline Patient Encoders

We compared our transformer-based encoder to multiple baseline encoders with varying degrees of complexity: (i) Average takes a simple average of features across cells, as is currently common practice when summarizing scRNA-seq to the patient level. (ii) Cell type fractions and (iii) Cell type means summarize cell type level information, either the fractions of cells in the sample assigned to each cell type, or the average of features for cells of a given specific cell type, concatenated together for all cell types in the dataset. (iv) Cell type fracs+means is the concatenation of the two. These cell type level summary vectors have been shown to correlate with clinical features (33; 44), but require expert labeling of cell types in order to construct, thus representing an expert-engineered baseline. (v) Kmeans cluster data into K clusters and concatenate the mean embeddings from each cluster. The K-means is trained on the training set, with results from K=30 and K=60 shown here. (vi) Inspired by recent work from (2), scSet w/ flow decoder uses a flow-based decoder during pretraining instead of a diffusion model. (vii) Finally, we compare to another attention-based encoder, ABMIL (16), as recent work for supervised clinical prediction from scRNA-seq employed this architecture (16; 11). Of note, while ABMIL uses attention to compute a parameterized weighted average of cells, it does not compute *self-attention* between input cells as our transformer encoder does.

To tease apart the benefit afforded by the architecture of the encoder vs. our diffusion pretraining, we included ablation baselines for each of the parametric encoders, evaluating the performance of scSet and ABMIL encoders both with and without diffusion pretraining.

#### 4.2.3. Prediction models

We input the patient embeddings to 3 different prediction models: (i) Linear probe, an L2-regularized logistic regression model, (ii) MLP, a simple multilayer perceptron with 2 hidden layers and GELU activations (14), and (iii) Finetune End-TO-End (FT-E2E), which uses the MLP from (ii) but jointly finetunes the encoder and MLP end-to-end, allowing gradient updates to propagate through both components to adapt the encoder’s representations for the downstream task.

For the MLP-based prediction heads, we use a weighted cross-entropy loss to compensate for class imbalance. Hyperparameter tuning was performed using nested K-fold validation, with inner K=5 and outer K=10, as described in Section 4.2.1. Hyperparameter details are provided in Appendix D.

#### 4.2.4. Metrics

We report the F1 score (↑) across folds, which balances precision and recall, making it suitable for class-imbalanced settings. For multiclass tasks, we use the weighted F1 score, averaging per-class F1 values weighted by the number of positives. Accuracy and AUC scores are reported in Appendix I.

#### 4.2.5. Results

Across most tasks and prediction models, scSet outperforms all other encoders and ablation models (Table 1). Our ablation baselines (SCSet w/o diffusion and ABMIL W/O DIFFUSION) suggest that pretraining the encoder via our conditional-diffusion autoencoder improves downstream supervised performance.

**Table 1.** Performance of scSet and baselines on clinical prediction tasks, as described in Section 4.2. Average weighted F1-Scores across folds *±* SEM are shown. Best performers (by mean) are bolded.

| TASK | COVID-19 |  |  | HLCA TRIPLE |  |  | SLE |  |  |
| --- | --- | --- | --- | --- | --- | --- | --- | --- | --- |
|  | LINEAR PROBE | MLP | FT-E2E | LINEAR PROBE | MLP | FT-E2E | LINEAR PROBE | MLP | FT-E2E |
| SCSET | **.95±.02** | **.93±.02** | **.93±.02** | .78±.06 | .68±.07 | **.66±.08** | .93±.02 | **.94±.01** | **0.95±0.01** |
| SCSET W/O DIFFUSION | .9±.03 | .86±.02 | .88±.03 | .61±.06 | .47±.08 | .53±.09 | .92±.02 | .83±.04 | .92±.02 |
| SCSET W/ FLOW DECODER | .9±.03 | .71±.04 | .87±.03 | .57±.07 | .43±.09 | .57±.07 | .87±.02 | .76±.04 | .92±.01 |
| ABMIL W/ DIFFUSION | .87±.03 | .83±.03 | .85±.03 | .62±.08 | .57±.06 | .52±.05 | .87±.02 | .9±.01 | .92±.01 |
| ABMIL W/O DIFFUSION | .88±.03 | .86±.03 | .84±.03 | .57±.07 | .38±.07 | .52±.06 | .87±.02 | .82±.01 | .9±.02 |
| AVERAGE | 0.88±.02 | .8±.04 | .81±.04 | .58±.07 | .49±.06 | .5±.05 | .88±.02 | .75±.03 | .77±.03 |
| CELL TYPE FRACTIONS | .84±.02 | .68±.04 | N/A | .6±.07 | .51±.09 | N/A | .83±.02 | .72±.03 | N/A |
| CELL TYPE MEANS | .87±.03 | .92±.03 | N/A | .73±.04 | .68±.04 | N/A | .94±.01 | .86±.03 | N/A |
| CELL TYPE FRACS+MEANS | .87±.03 | .92±.03 | N/A | .72±.04 | **.7±.05** | N/A | **.95±.01** | .9±.02 | N/A |
| KMEANS30 | .92±.03 | .86±.04 | N/A | .77±.05 | .63±.06 | N/A | .94±.02 | .92±.02 | N/A |
| KMEANS60 | .94±.02 | .9±.02 | N/A | **.79±.04** | .66±.06 | N/A | .89±.02 | .92±.01 | N/A |

In real-world settings, clinically-labeled scRNA-seq cohorts are often small (3), and thus a model that can improve predictive performance on small amounts of labeled data is valuable. With this in mind, we evaluated each model’s performance when trained on just 25, 50, or 100 training samples per-fold. We repeated this experiment five times, each time using a different random subset of data, and we report the mean and standard error of the mean (SEM) across all random subsets and test folds. Even with limited training data, scSet consistently outperforms the baseline models (Figure 6).

**Figure 6.**
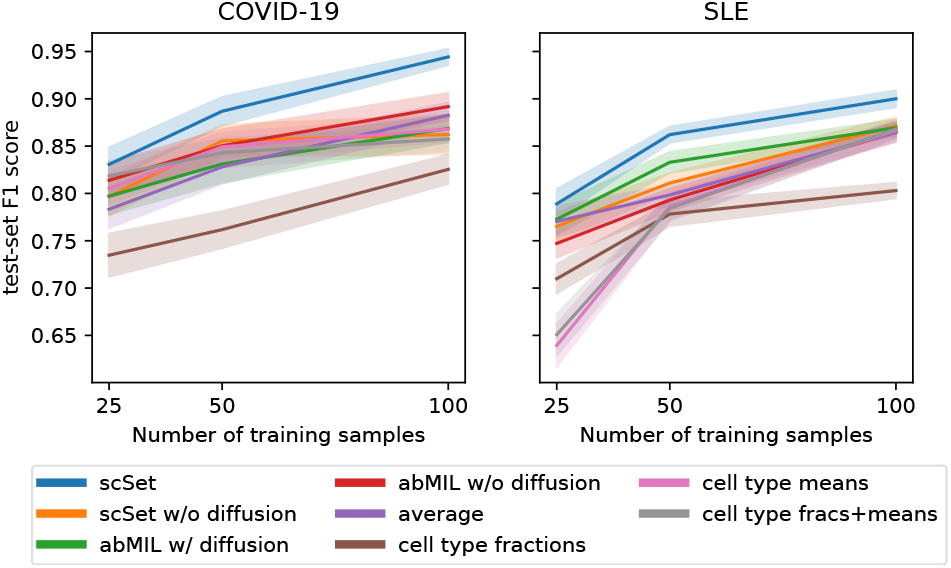
With limited numbers of training samples, scSet still outperforms baseline encoders on the COVID-19 and SLE prediction tasks. Error bars represent the standard error, calculated over 5 random subsamplings of the training data for each of 10 folds.

## 5. Discussion

Our results demonstrate that scSet effectively learns meaningful patient-level representations from single-cell RNA sequencing data through a diffusion-based autoencoding framework. By leveraging a transformer-based encoder to aggregate unordered single-cells, and employing a conditional diffusion decoder to generate realistic cellular compositions, scSet provides a powerful and flexible method for patient-level modeling that elegantly circumvents the challenge of autoencoding set-structured data. scSet embeddings prove useful for downstream clinical prediction tasks, suggesting that scSet captures clinically relevant signals that generalize across datasets.

The introduction of self-supervised learning for patient-level representations from scRNA-seq data mitigates the common issue of limited labeled datasets in biomedical applications. By learning from large-scale unlabeled data, scSet can create pretrained representations that transfer effectively to new clinical prediction tasks with minimal labeled data. This approach is particularly advantageous for studying rare diseases or heterogeneous conditions where labeled single-cell samples are scarce. Additionally, our framework is modular and can incorporate different cell embeddings, making it adaptable to future advances in single-cell representation learning.

## 6. Limitations and Future Work

While scSet presents a promising framework for patient-level representation learning, several limitations remain. First, our model is trained on cells represented by precomputed scVI embeddings. While this standardizes inputs and mitigates batch effects, future work could explore end-to-end training of cell and patient embeddings to improve interpretability and performance.

Additionally, our current approach conditions the generative diffusion model on patient representations derived from scRNA-seq data; however, this framework could be extended to generate single-cell profiles conditioned on bulk RNA-seq profiles, patient characteristics, or single-cell data from other modalities.

As spatial scRNA-seq data becomes more widely available, future extensions could integrate spatial information, which is often predictive of clinical outcomes (41; 35).

Importantly, although this work aims to bridge single-cell data and translational impact, it remains early-stage and would require further development, validation, and clinical testing before clinical use.

## Acknowledgements

We thank Nalini Singh for collaborating on early iterations of this work and Marinka Zitnik for her feedback on experimental design and interpretation. R.B., J.W., and D.S were supported by an ASPIRE award from The Mark Foundation for Cancer Research. J.W. was additionally supported by the Swiss Study Foundation. G.G. is partially supported by the Paul C. Zamecnik Chair in Oncology at the Massachusetts General Hospital Cancer Center.

## Technical Appendices and Supplementary Material

### A. Interpretation of scSet as an Autoregressive Model

Our work shares a conceptual connection with autoregressive modeling. Viewing each cell in a sample as a token similar to those used in natural language processing, scSet aims to learn the joint distribution over tokens, for which autoregressive approaches have recently shown great capabilities (5). However, autoregressive models typically process sequences, where there is a canonical ordering that determines the next token to predict. Further, autoregressive approaches generally learn a probability distribution over discrete tokens, not continuous samples such as single-cell profiles. Inspired by the framework introduced in Li et al. (2024) (20), scSet can be seen as an autoregressive model which selects an arbitrary ordering of cells for each sample, and then uses the first *N*_anchor_ *< N* cells to predict a latent prototype of the next cell state, namely the patient representation, which is subsequently transformed into an actual cell state by the diffusion model.

### B. Data Pre-Processing

The scSet autoencoder was trained on data from the CZ CELLxGENE (CxG) Discover Census (3), which we filtered to the 7,342 samples with at least 128 cells.

### C. Algorithm for Pretraining scSet

#### Algorithm 1

Conditional Diffusion Autoencoding Training

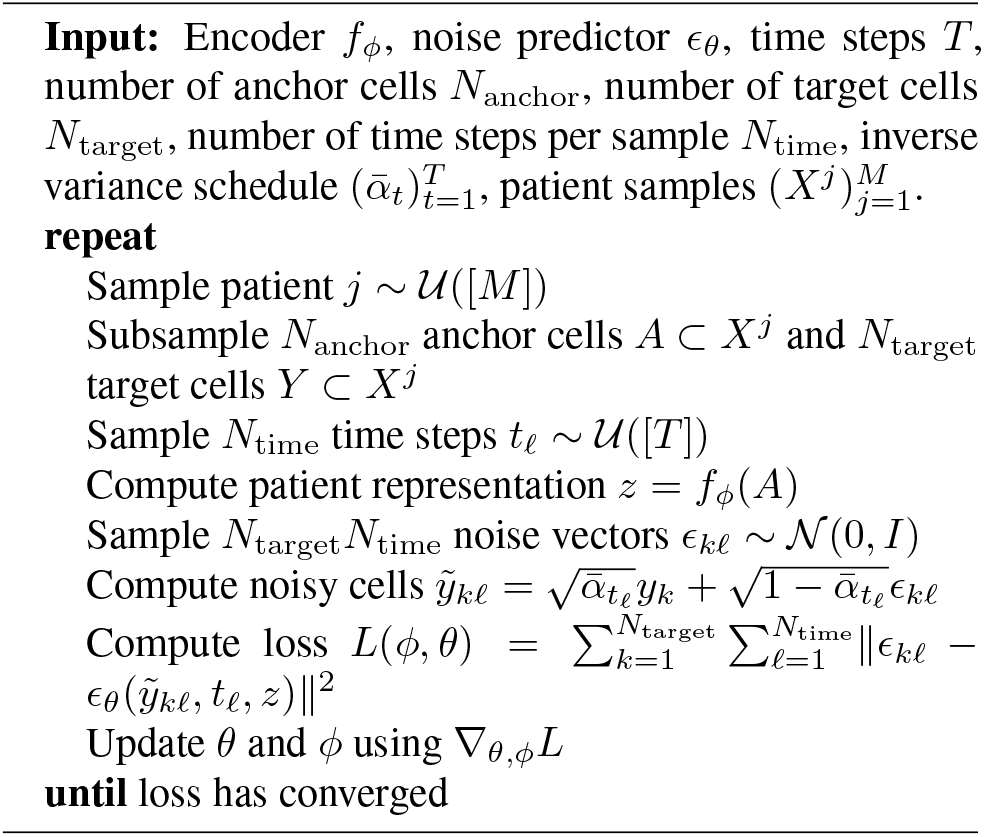

### D. Hyperparameter Tuning

For our scset model, which is made up of a transformer and a denoising diffusion network, we used the following hyperparameters for the transformer: 4 transformer heads, 2 blocks (layers) of transformers, batch size 32, and learning rate 10^−3^. We searched over the following hyperparameters: number transformer heads {2, 4} ; number transformer blocks {2, 3, 4} ; batch size {16, 32, 64, 128} ; learning rate {10^−2^, 10^−3^, 10^−4^}. We found that our choice of batch size and learning rate significantly affected validation set denoising loss. While models trained with different batch sizes converged to a similar loss by 200 epochs, we found that the larger the batch size, the longer the model took to converge to this loss. We chose a batch size of 32, to balance this behavior (which would suggest choosing the lowest batch size) with efficient use of our GPUs (which are only partially utilized at lower batch sizes). Once the batch size was fixed, a learning rate of 10^−3^ performed best. The number of transformer heads and blocks did not meaningfully alter performance, so we settled on 4 heads and 2 blocks to balance expressivity with avoiding overfitting and unnecessary complexity.

For the logistic regression in sklearn, we tuned the hyperparameter C over the following values using nested cross-validation: [0.01, 0.1, 1, 10, 100, 1000, 10000, 100000, 1000000].

### E. Compute Environment

Models were trained on a single NVIDIA A100 80GB GPU, on a cluster with 504GB RAM.

### F. Statistics

We ran each task on K held out test folds. We report the 95% confidence intervals for the mean performance across these folds. To calculate these intervals, we determined the sample mean (x) and sample standard deviation (s) for the performance metrics, then computed the standard error of the mean (SEM) as 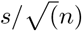, where *n* is the number of runs (*n* = 10). For *n* = 10, the t-value for a 95% confidence level is 2.262. The margin of error (ME) was obtained by multiplying the t-value with the SEM. We reported 95% confidence intervals as x*±*ME.

### G. Details for Generating Semi-Synthetic Datasets

scVI embeddings for cells from a multiple myeloma study (43) were obtained using the trained scVI model downloaded from CZ CELLxGENE Discover Census (3) used throughout this work.

To create the semi-synthetic data for our perturbation subtyping experiment, we first created 12 patients with 200 cells each, sampling 25% of cells from each cell type (natural killer cells, helper T cells, CD8+ cytotoxic T cells, and monocytes). For the 6 patients assigned to our ‘perturbed’ subtype, we added a constant to five of the latent dimensions for all helper T cells. We subtracted this same constant from the same latents for all CD8+ cytotoxic T cells in those sample samples. We chose this perturbation structure because it represents a case where averaging, or pseudobulking, the sample would lose the signal due to the equal and opposite effect of the two perturbations.

### H. Tables Comparing Cell Type Proportions Between True and scSet Reconstructed Samples

See Tables 2, 3, 4, 5.

**Table 2.**
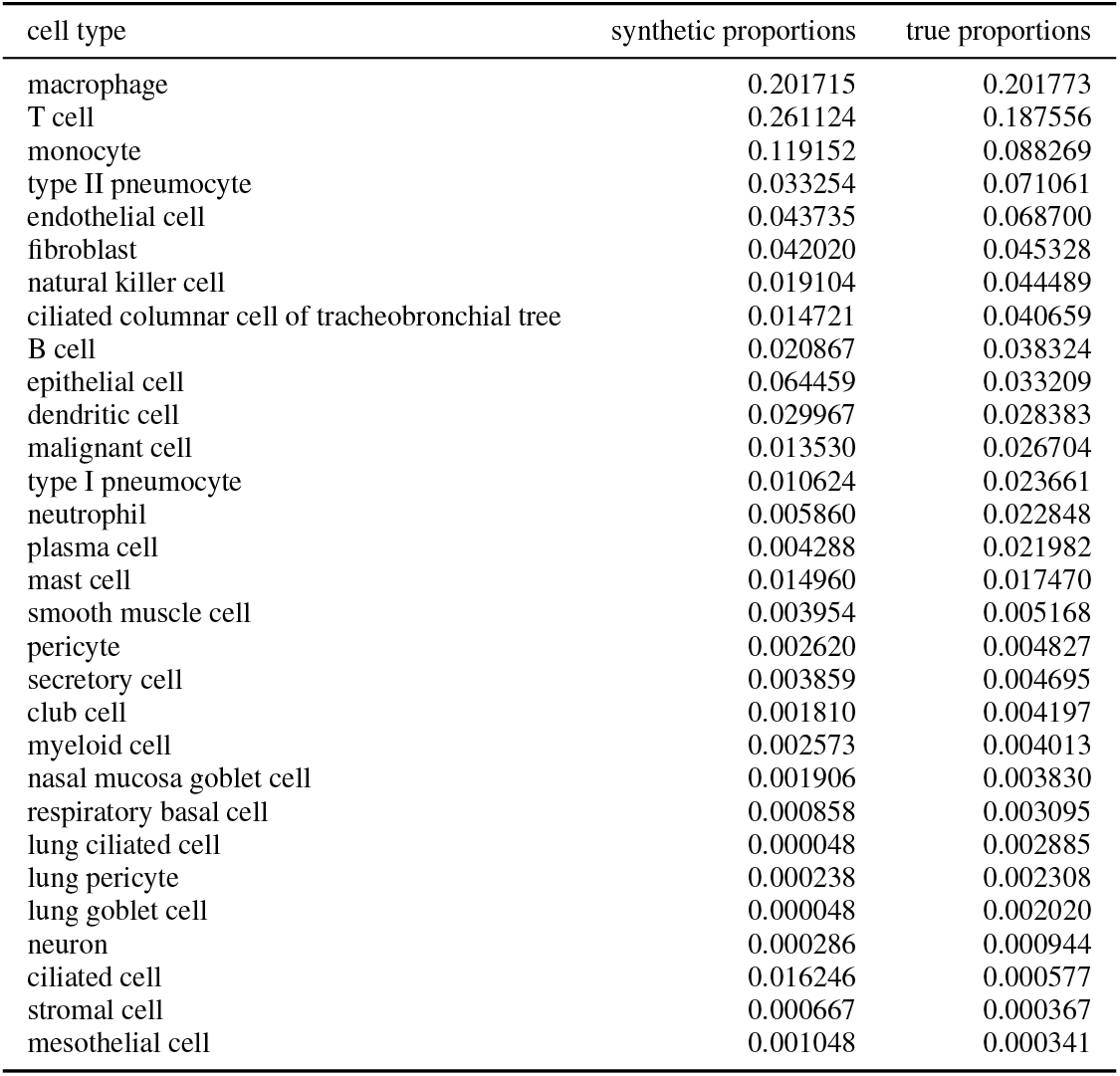
Cell type proportions for true and scSet reconstructed **lung** samples. Limited to 30 most common cell types among true cells.

| cell type | synthetic proportions | true proportions |
| --- | --- | --- |
| macrophage | 0.201715 | 0.201773 |
| T cell | 0.261124 | 0.187556 |
| monocyte | 0.119152 | 0.088269 |
| type II pneumocyte | 0.033254 | 0.071061 |
| endothelial cell | 0.043735 | 0.068700 |
| fibroblast | 0.042020 | 0.045328 |
| natural killer cell | 0.019104 | 0.044489 |
| ciliated columnar cell of tracheobronchial tree | 0.014721 | 0.040659 |
| B cell | 0.020867 | 0.038324 |
| epithelial cell | 0.064459 | 0.033209 |
| dendritic cell | 0.029967 | 0.028383 |
| malignant cell | 0.013530 | 0.026704 |
| type I pneumocyte | 0.010624 | 0.023661 |
| neutrophil | 0.005860 | 0.022848 |
| plasma cell | 0.004288 | 0.021982 |
| mast cell | 0.014960 | 0.017470 |
| smooth muscle cell | 0.003954 | 0.005168 |
| pericyte | 0.002620 | 0.004827 |
| secretory cell | 0.003859 | 0.004695 |
| club cell | 0.001810 | 0.004197 |
| myeloid cell | 0.002573 | 0.004013 |
| nasal mucosa goblet cell | 0.001906 | 0.003830 |
| respiratory basal cell | 0.000858 | 0.003095 |
| lung ciliated cell | 0.000048 | 0.002885 |
| lung pericyte | 0.000238 | 0.002308 |
| lung goblet cell | 0.000048 | 0.002020 |
| neuron | 0.000286 | 0.000944 |
| ciliated cell | 0.016246 | 0.000577 |
| stromal cell | 0.000667 | 0.000367 |
| mesothelial cell | 0.001048 | 0.000341 |

**Table 3.** Cell type proportions for true and scSet reconstructed **blood** samples. Limited to 30 most common cell types among true cells.

| cell type | synthetic proportions | true proportions |
| --- | --- | --- |
| T cell | 0.671453 | 0.407475 |
| naive T cell | 0.017003 | 0.168529 |
| monocyte | 0.161800 | 0.156635 |
| natural killer cell | 0.030631 | 0.122473 |
| B cell | 0.101011 | 0.064810 |
| naive B cell | 0.003179 | 0.040356 |
| dendritic cell | 0.011306 | 0.011751 |
| platelet | 0.000768 | 0.006774 |
| plasmablast | 0.000804 | 0.003751 |
| T-helper | 0.000009 | 0.003279 |
| blood cell | 0.000357 | 0.003261 |
| T follicular helper cell | 0.000071 | 0.002978 |
| progenitor cell | 0.000911 | 0.001455 |
| erythrocyte | 0.000170 | 0.001065 |
| lymphocyte | 0.000143 | 0.001038 |
| thymocyte | 0.000054 | 0.000916 |
| plasma cell | 0.000036 | 0.000687 |
| IgG plasma cell | 0.000009 | 0.000445 |
| IgA plasma cell | 0.000009 | 0.000427 |
| double negative T regulatory cell | 0.000009 | 0.000337 |
| innate lymphoid cell | 0.000036 | 0.000296 |
| macrophage | 0.000054 | 0.000225 |
| IgA plasmablast | 0.000161 | 0.000180 |
| common lymphoid progenitor | 0.000009 | 0.000180 |
| megakaryocyte | 0.000000 | 0.000166 |
| ILC1, human | 0.000000 | 0.000126 |
| myeloid cell | 0.000000 | 0.000117 |
| IgM plasma cell | 0.000000 | 0.000108 |
| IgG plasmablast | 0.000009 | 0.000085 |
| megakaryocyte-erythroid progenitor cell | 0.000000 | 0.000076 |

**Table 4.** Cell type proportions for true and scSet reconstructed **breast** samples. Limited to 30 most common cell types among true cells.

| cell type | synthetic proportions | true proportions |
| --- | --- | --- |
| epithelial cell | 0.270300 | 0.204534 |
| fibroblast | 0.139667 | 0.146364 |
| T cell | 0.158140 | 0.115783 |
| endothelial cell | 0.073285 | 0.092961 |
| progenitor cell | 0.079273 | 0.089614 |
| basal cell | 0.070646 | 0.079420 |
| endothelial tip cell | 0.070443 | 0.060402 |
| luminal hormone-sensing cell of mammary gland | 0.024259 | 0.046506 |
| perivascular cell | 0.017052 | 0.036515 |
| pericyte | 0.009541 | 0.022112 |
| macrophage | 0.046488 | 0.021452 |
| smooth muscle cell | 0.002436 | 0.018359 |
| subcutaneous adipocyte | 0.007917 | 0.014657 |
| B cell | 0.006293 | 0.008317 |
| plasmablast | 0.001827 | 0.007151 |
| naive B cell | 0.000812 | 0.005376 |
| natural killer cell | 0.001827 | 0.004970 |
| monocyte | 0.007714 | 0.004564 |
| IgA plasma cell | 0.000812 | 0.004311 |
| dendritic cell | 0.006192 | 0.003398 |
| naive T cell | 0.000000 | 0.002891 |
| lymphocyte | 0.000000 | 0.002434 |
| myeloid cell | 0.001320 | 0.002384 |
| Tc1 cell | 0.000000 | 0.001471 |
| leukocyte | 0.000000 | 0.001065 |
| mast cell | 0.002944 | 0.001014 |
| contractile cell | 0.000406 | 0.000710 |
| IgG plasma cell | 0.000203 | 0.000710 |
| neutrophil | 0.000203 | 0.000558 |

**Table 5.**
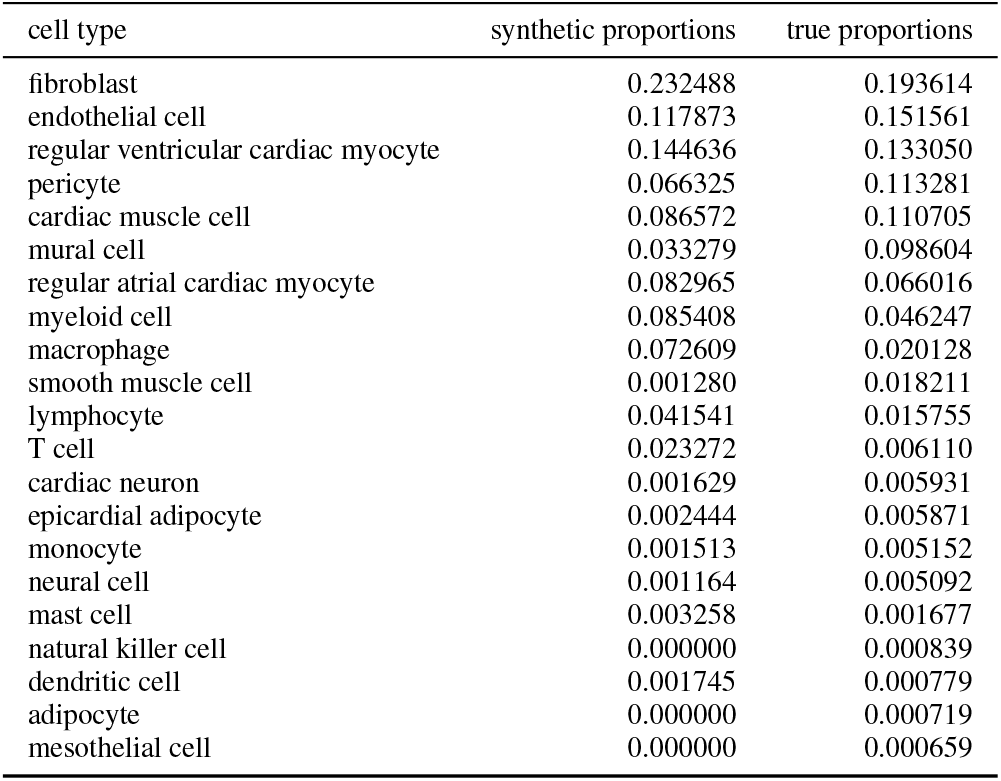
Cell type proportions for true and scSet reconstructed **heart** samples. Limited to 30 most common cell types among true cells.

| cell type | synthetic proportions | true proportions |
| --- | --- | --- |
| fibroblast | 0.232488 | 0.193614 |
| endothelial cell | 0.117873 | 0.151561 |
| regular ventricular cardiac myocyte | 0.144636 | 0.133050 |
| pericyte | 0.066325 | 0.113281 |
| cardiac muscle cell | 0.086572 | 0.110705 |
| mural cell | 0.033279 | 0.098604 |
| regular atrial cardiac myocyte | 0.082965 | 0.066016 |
| myeloid cell | 0.085408 | 0.046247 |
| macrophage | 0.072609 | 0.020128 |
| smooth muscle cell | 0.001280 | 0.018211 |
| lymphocyte | 0.041541 | 0.015755 |
| T cell | 0.023272 | 0.006110 |
| cardiac neuron | 0.001629 | 0.005931 |
| epicardial adipocyte | 0.002444 | 0.005871 |
| monocyte | 0.001513 | 0.005152 |
| neural cell | 0.001164 | 0.005092 |
| mast cell | 0.003258 | 0.001677 |
| natural killer cell | 0.000000 | 0.000839 |
| dendritic cell | 0.001745 | 0.000779 |
| adipocyte | 0.000000 | 0.000719 |
| mesothelial cell | 0.000000 | 0.000659 |

As described in Section 4.1.2, we expect scSet’s diffusion decoder to generate sets of cells that closely resemble each patient’s true set of cells. Thus, we evaluated scSet’s reconstruction capabilities using patient samples from the test set: cells from each test set patient were encoded to a patient embedding using the trained encoder, *z*_*j*_ = *f*_*θ*_(*X*^*j*^). We then generated 500 cells per patient (the number of decoded cells is arbitrarily set by the user). Starting from Gaussian noise at time step *T* = 1000, we visualize the UMAPs (27) of the reconstructed cell profiles of multiple patients grouped by tissue as they are denoised over time steps in Figure 2. We found that for each tissue, the model generates the expected cell types in relatively correct proportions. Here, we include cell type proportions tables to give more insight into this result. For each tissue shown in Figure 2, we include a table of the true and simulated cell type compositions for each tissue (this data is aggregated across samples from each tissue, i.e. we took the union of cells from all samples belonging to each tissue). The true and simulated proportions are relatively close, which can be summarized by the Pearson correlation coefficient between the two columns in each table, which is consistently high: 0.95 for lung, 0.97 for breast, 0.89 for heart, and 0.91 for blood. Note that since generated cells have no ground truth cell type labels, we predicted instead pseudo-cell type labels from their simulated profiles at *t* = 0 using a *k*-Nearest Neighbors classifier trained on the true cells from these patients.

### I. Full Results for All Clinical Prediction Tasks

See Tables 6, 7, 8, 9.

**Table 6.** Full set of results for the triple HLCA task. Average across folds *±* SEM are shown.

| MODEL | AUC |  |  | ACCURACY |  |  | WEIGHTED F1 |  |  |
| --- | --- | --- | --- | --- | --- | --- | --- | --- | --- |
|  | LINEAR PROBE | MLP | FT-E2E | LINEAR PROBE | MLP | FT-E2E | LINEAR PROBE | MLP | FT-E2E |
| SCSet | 0.89±0.05 | 0.83±0.06 | 0.74±0.11 | 0.75±0.06 | 0.66±0.07 | 0.65±0.07 | 0.78±0.06 | 0.68±0.07 | 0.66±0.08 |
| SCSet w/o DIFFUSION | 0.73±0.07 | 0.71±0.05 | 0.72±0.07 | 0.61±0.05 | 0.45±0.07 | 0.51±0.08 | 0.61±0.06 | 0.47±0.08 | 0.53±0.09 |
| SCSet w/ FLOW DECODER | 0.73±0.07 | 0.66±0.06 | 0.73±0.07 | 0.58±0.06 | 0.49±0.09 | 0.55±0.07 | 0.57±0.07 | 0.43±0.09 | 0.57±0.07 |
| ABMIL w/ DIFFUSION | 0.71±0.07 | 0.7±0.05 | 0.71±0.05 | 0.65±0.08 | 0.53±0.06 | 0.48±0.06 | 0.62±0.08 | 0.57±0.06 | 0.52±0.05 |
| ABMIL w/o DIFFUSION | 0.71±0.07 | 0.68±0.05 | 0.73±0.05 | 0.58±0.06 | 0.39±0.06 | 0.49±0.06 | 0.57±0.07 | 0.38±0.07 | 0.52±0.06 |
| AVERAGE | 0.71±0.06 | 0.62±0.05 | 0.64±0.05 | 0.62±0.07 | 0.44±0.05 | 0.46±0.04 | 0.58±0.07 | 0.49±0.06 | 0.5±0.05 |
| CELL TYPE FRACTIONS | 0.79±0.04 | 0.74±0.1 | NAN | 0.58±0.06 | 0.47±0.09 | NAN | 0.6±0.07 | 0.51±0.09 | NAN |
| CELL TYPE MEANS | 0.9±0.03 | 0.84±0.05 | NAN | 0.7±0.04 | 0.63±0.04 | NAN | 0.73±0.04 | 0.68±0.04 | NAN |
| CELL TYPE FRACS+MEANS | 0.9±0.03 | 0.8±0.06 | NAN | 0.7±0.04 | 0.66±0.05 | NAN | 0.72±0.04 | 0.7±0.05 | NAN |
| KMEANS30 | 0.92±0.05 | 0.83±0.08 | NAN | 0.76±0.05 | 0.58±0.07 | NAN | 0.77±0.05 | 0.63±0.06 | NAN |
| KMEANS60 | 0.9±0.03 | 0.79±0.07 | NAN | 0.78±0.04 | 0.63±0.07 | NAN | 0.79±0.04 | 0.66±0.06 | NAN |

**Table 7.**
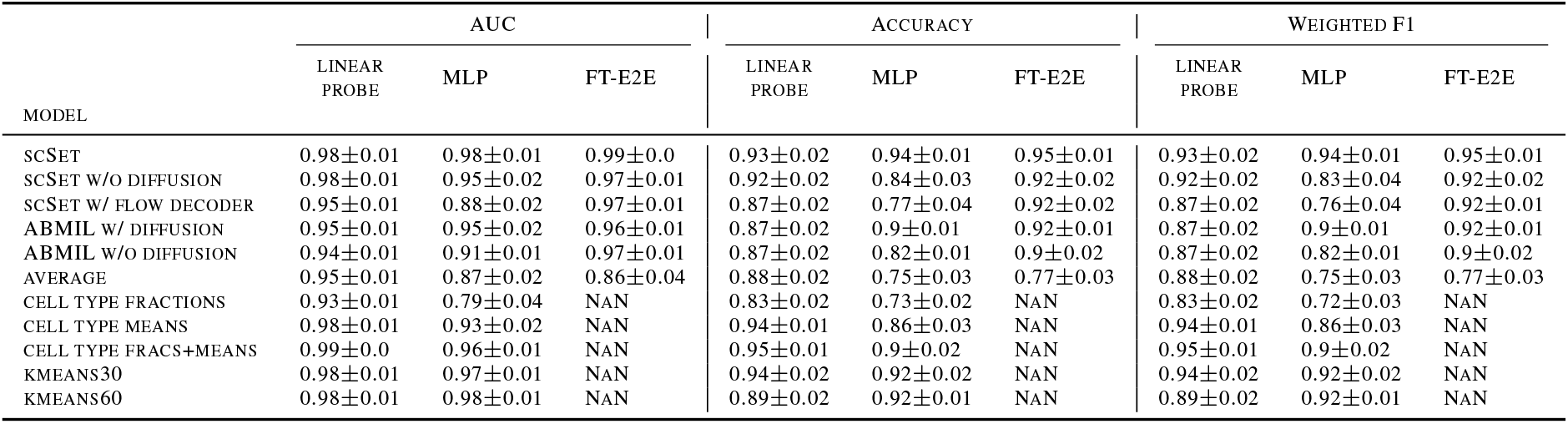
Full set of results for the SLE task. Average across folds *±* SEM are shown.

**Table 8.** Full set of results for the COVID-19 task. Average across folds *±* SEM are shown.

| MODEL | AUC |  |  | ACCURACY |  |  | WEIGHTED F1 |  |  |
| --- | --- | --- | --- | --- | --- | --- | --- | --- | --- |
|  | LINEAR PROBE | MLP | FT-E2E | LINEAR PROBE | MLP | FT-E2E | LINEAR PROBE | MLP | FT-E2E |
| SCSet | 0.98±0.01 | 0.98±0.02 | 0.98±0.01 | 0.95±0.02 | 0.93±0.02 | 0.92±0.02 | 0.95±0.02 | 0.93±0.02 | 0.93±0.02 |
| SCSet w/o DIFFUSION | 0.95±0.02 | 0.96±0.02 | 0.82±0.1 | 0.9±0.02 | 0.86±0.02 | 0.88±0.03 | 0.9±0.03 | 0.86±0.02 | 0.88±0.03 |
| SCSet w/ FLOW DECODER | 0.93±0.02 | 0.79±0.04 | 0.94±0.05 | 0.89±0.03 | 0.75±0.02 | 0.86±0.03 | 0.9±0.03 | 0.71±0.04 | 0.87±0.03 |
| ABMIL w/ DIFFUSION | 0.92±0.03 | 0.86±0.06 | 0.88±0.06 | 0.88±0.03 | 0.83±0.03 | 0.85±0.03 | 0.87±0.03 | 0.83±0.03 | 0.85±0.03 |
| ABMIL w/o DIFFUSION | 0.93±0.03 | 0.92±0.03 | 0.9±0.04 | 0.89±0.03 | 0.85±0.03 | 0.86±0.02 | 0.88±0.03 | 0.86±0.03 | 0.84±0.03 |
| AVERAGE | 0.94±0.02 | 0.85±0.03 | 0.86±0.04 | 0.88±0.02 | 0.79±0.04 | 0.79±0.05 | 0.88±0.02 | 0.8±0.04 | 0.81±0.04 |
| CELL TYPE FRACTIONS | 0.89±0.04 | 0.61±0.06 | NAN | 0.85±0.02 | 0.67±0.04 | NAN | 0.84±0.02 | 0.68±0.04 | NAN |
| CELL TYPE MEANS | 0.96±0.02 | 0.97±0.02 | NAN | 0.88±0.02 | 0.93±0.03 | NAN | 0.87±0.03 | 0.92±0.03 | NAN |
| CELL TYPE FRACS+MEANS | 0.96±0.02 | 0.97±0.02 | NAN | 0.88±0.02 | 0.91±0.03 | NAN | 0.87±0.03 | 0.92±0.03 | NAN |
| KMEANS30 | 0.95±0.02 | 0.9±0.04 | NAN | 0.92±0.02 | 0.87±0.04 | NAN | 0.92±0.03 | 0.86±0.04 | NAN |
| KMEANS60 | 0.96±0.01 | 0.97±0.02 | NAN | 0.94±0.02 | 0.9±0.02 | NAN | 0.94±0.02 | 0.9±0.02 | NAN |

**Table 9.** Full set of results for the binary HLCA task. Average across folds *±* SEM are shown.

| MODEL | AUC |  |  | ACCURACY |  |  | WEIGHTED F1 |  |  |
| --- | --- | --- | --- | --- | --- | --- | --- | --- | --- |
|  | LINEAR PROBE | MLP | FT-E2E | LINEAR PROBE | MLP | FT-E2E | LINEAR PROBE | MLP | FT-E2E |
| SCSet | 0.78±0.06 | 0.81±0.06 | 0.85±0.03 | 0.78±0.06 | 0.87±0.04 | 0.83±0.06 | 0.76±0.09 | 0.87±0.04 | 0.83±0.06 |
| SCSet w/o DIFFUSION | 0.64±0.09 | 0.58±0.1 | 0.71±0.08 | 0.8±0.07 | 0.71±0.07 | 0.81±0.06 | 0.77±0.09 | 0.73±0.06 | 0.82±0.06 |
| SCSet w/ FLOW DECODER | 0.47±0.09 | 0.48±0.15 | 0.73±0.04 | 0.81±0.07 | 0.72±0.1 | 0.84±0.04 | 0.75±0.09 | 0.67±0.11 | 0.85±0.04 |
| ABMIL w/ DIFFUSION | 0.59±0.06 | 0.66±0.08 | 0.68±0.06 | 0.79±0.07 | 0.77±0.07 | 0.79±0.05 | 0.74±0.09 | 0.78±0.07 | 0.78±0.06 |
| ABMIL w/o DIFFUSION | 0.46±0.13 | 0.58±0.11 | 0.45±0.09 | 0.81±0.07 | 0.7±0.07 | 0.7±0.07 | 0.75±0.09 | 0.7±0.08 | 0.7±0.07 |
| AVERAGE | 0.51±0.12 | 0.64±0.12 | 0.48±0.06 | 0.81±0.07 | 0.69±0.06 | 0.63±0.04 | 0.75±0.09 | 0.73±0.06 | 0.67±0.05 |
| CELL TYPE FRACTIONS | 0.81±0.07 | 0.53±0.08 | NAN | 0.79±0.07 | 0.7±0.06 | NAN | 0.77±0.09 | 0.73±0.07 | NAN |
| CELL TYPE MEANS | 0.9±0.07 | 0.82±0.06 | NAN | 0.86±0.04 | 0.72±0.07 | NAN | 0.85±0.05 | 0.76±0.06 | NAN |
| CELL TYPE FRACS+MEANS | 0.9±0.06 | 0.81±0.08 | NAN | 0.86±0.04 | 0.79±0.04 | NAN | 0.85±0.05 | 0.82±0.04 | NAN |
| KMEANS30 | 0.9±0.04 | 0.82±0.04 | NAN | 0.88±0.03 | 0.87±0.04 | NAN | 0.89±0.03 | 0.87±0.04 | NAN |
| KMEANS60 | 0.92±0.04 | 0.86±0.03 | NAN | 0.87±0.04 | 0.82±0.03 | NAN | 0.86±0.04 | 0.83±0.04 | NAN |

